# Cardiopulmonary structural, functional and immune-alterations in a Down syndrome mouse model and upon modulation of EGCG

**DOI:** 10.1101/2023.03.13.532396

**Authors:** Birger Tielemans, Sergi Llambrich, Laura Seldeslachts, Jonathan Cremer, Hung Chang Tsui, Anne-Charlotte Jonckheere, Fopke Marain, Mirko Riedel, Jens Wouters, Julia Herzen, Bartosz Leszczyński, Erik Verbeken, Jeroen Vanoirbeek, Greetje Vande Velde

## Abstract

In individuals with Down syndrome (DS), cardiovascular and pulmonary diseases are the most common health problem and result in increased mortality and morbidity. Although these clinical comorbidities are well described, no preclinical models for DS are fully characterized for cardiopulmonary alterations, preventing research to understanding the development and pharmacological modulation of lungs, heart and immune system. Our objective is to characterize the cardiopulmonary and immunological phenotype in Ts65Dn mice and investigate the modulatory effects green tea extract enriched in epigallocatechin 3 gallate (GTE-EGCG). GTE-EGCG administration started at embryonic day 9 and was discontinued at postnatal day (PD) 180. Newborns were longitudinally monitored until PD210 using micro-computed tomography. At endpoint, we characterized the structural, functional and immunological alterations and persistent effects of GTE-EGCG administration. This study revealed normal lung development in the Ts65Dn mice and highlighted RV hypertrophy and immunological alterations. GTE-EGCG administration resulted in genotype-specific and genotype-independent alterations resulting in lung immaturation and airway hyperreactivity. Our results highlight the cardiovascular and immunological phenotype of Ts65Dn mice and potential use for safety studies of therapeutic agents in a DS-specific context.

**Summary statement:** This study longitudinally follows respiratory and cardiac alterations in the Ts65Dn mouse model and describes the impact of prenatal EGCG modulation on the euploid and trisomic phenotype

## Introduction

Down syndrome (DS) is the most common genetic cause of intellectual disability (Potier and Reeves, 2016). Apart from the neurological and cognitive deficits, individuals with DS present with a spectrum of abnormalities affecting most systems and body organs (Frid et al., 1999). As such, individuals with DS are more prone to develop various cardiopulmonary disorders including impaired pulmonary vascular growth leading to pulmonary hypoplasia and resulting in an increased incidence of pulmonary hypertension (Bush et al., 2018, 2017), presence of congenital heart diseases (CHD) with persistent left-to-right shunts, chronic upper airway obstruction (Benhaourech et al., 2016; Maris et al., 2016) and an impaired immune system highlighted by increased interleukin concentrations leading to a high incidence of leukemia (Khan et al., 2011; Zhang et al., 2017) and respiratory infections (Bruijn et al., 2007). Alterations in lung physiology include structural as well as functional and immunological alterations, resulting in mortality and morbidity in 75% of individuals with DS (Colvin and Yeager, 2017). Nevertheless, the majority of clinical and pre-clinical research towards DS is focussed on brain and neurological development, Alzheimer’s disease and cognitive deficits. Although the association between trisomy 21 and its whole-body comorbidities is clinically well described in terms of aetiology, pre-clinical studies towards the molecular background, lung pathophysiology and development of these respiratory, cardiopulmonary and underlying immune deficits in DS are lacking. Filling in this knowledge gap, i.e. understanding the pathophysiological development of cardiopulmonary and respiratory disorders in a fully characterized preclinical animal model, may help to find new prevention strategies and improve quality of life and life expectancy in individuals with DS.

Mouse orthologs of the human chromosome 21 (Hsa21) genes are spread across three mouse chromosomes (Mmu16, Mmu17 and Mmu10) making it challenging to resemble the complete human DS genotype and corresponding phenotype. Nevertheless, several mouse models exist that closely mimic the human DS phenotype in some or several aspects. One of the most commonly used preclinical animal models in DS research is the Ts65Dn, which contains triplication of the DS chromosomal region (Crété et al., 1993; Korenberg et al., 1997), also called DS critical region, a region of genes which is involved in all forms of human DS (Belichenko et al., 2009; Jiang et al., 2015; Olson et al., 2004). This model is very well characterized regarding the main hallmarks of DS including craniofacial malformations, brain dysmorphology, altered bone micro-architecture and cognitive impairment (Herault et al., 2017; Richtsmeier et al., 2000). However, a proper characterization and a (holistic) view on the pulmonary, cardiac and immunological development, studied in a DS-specific preclinical setting, is missing.

Nevertheless, pharmaceutical treatment options have already been explored, focussing entirely on cognitive and neuronal improvements. Between 2012 and 2014, a phase II clinical trial (TESDAD) investigated the safety and efficacy of green tea extracts containing epigallocatechin-3-gallate (EGCG) on young adults with DS as promising therapeutic tool and showed improvement in cognition (de la Torre et al., 2016). Ever since, the effect of green tea extracts (GTE) enriched in catechins such as EGCG and its metabolites is extensively studied towards its effect on cognition, neurological and craniofacial development (Catuara-Solarz et al., 2015; De la Torre et al., 2014; Stagni et al., 2015; Starbuck et al., 2021). EGCG, as most abundant and active catechin in GTE, was demonstrated to also have antiatherogenic and antioxidative properties suggesting a potential protective effect on the cardiovascular system (Hayek et al., 1997; Ludwig et al., 2004; Miura et al., 2000; Yang and Koo, 2000). EGCG is a natural inhibitor of the protein dual-specificity tyrosine-phosphorylation regulated protein kinase 1A (*DYRK1A*), which is located in the DS critical region and thus overexpressed in DS (De la Torre et al., 2014). Overexpression of DYRK1A is a key player in a complex clinical syndrome including defective pulmonary vascular development affecting important factors including vascular endothelial growth factor (VEGF) and repressing activity of nuclear factor of activated T-cells (NFATc) (Fernández-Martínez et al., 2015; Rozen et al., 2018). However, the effects of pharmacological modulation using GTE-EGCG on the lung, heart and immune system in DS have never been assessed, which warrants an integrated multi-system evaluation towards safety and dose regimen since potential side effects during administration of GTE-EGCG for cognitive improvements are currently overlooked.

Therefore, In the present study, we (1) characterized development of the lung and heart phenotype in the Ts65Dn mouse model of DS together with the immunological profile at adulthood by focussing on structural, functional and immunological alterations compared to WT littermates. We performed longitudinal *in vivo* micro-computed tomography (µCT) from birth until adulthood to characterize the macroscopic development of the lungs. We complemented our findings with echocardiography and contrast-enhanced µCT adulthood and performed lung function analysis at endpoint. In addition, microscopic analysis of heart, lung and immune alterations were conducted by histopathology, phase contrast CT and flow cytometry at endpoint; (2) we investigated the impact of perinatal GTE-EGCG administration as DS modulating agent on the cardiopulmonary development of both wild type and Ts65Dn mice and follow-up on the persistent effects of GTE-EGCG after treatment discontinuation.

Therefore, this study provides important missing data on the characterization of the Ts65Dn mouse model in terms of their cardiopulmonary and immunological phenotype and describe the effect of GTE-EGCG on the heart, lungs and immune profile in euploid and trisomic mice, providing essential information on safe GTE-EGCG supplementation.

## Results

### Characterization of the structural and functional pulmonary phenotype of Ts65Dn mice throughout development and modulatory effects of GTE-EGCG

We first characterized the structural and functional pulmonary phenotype in WT and TS mice together with the modulatory effects of GTE-EGCG treatment throughout development by means of micro-computed tomography (µCT) and histology and phase-contrast CT at endpoint. At endpoint we performed lung function testing and histopathology to obtain structural and functional information on the lung in Ts65Dn and wild type mice (Figure 1A).

**Figure 1:**
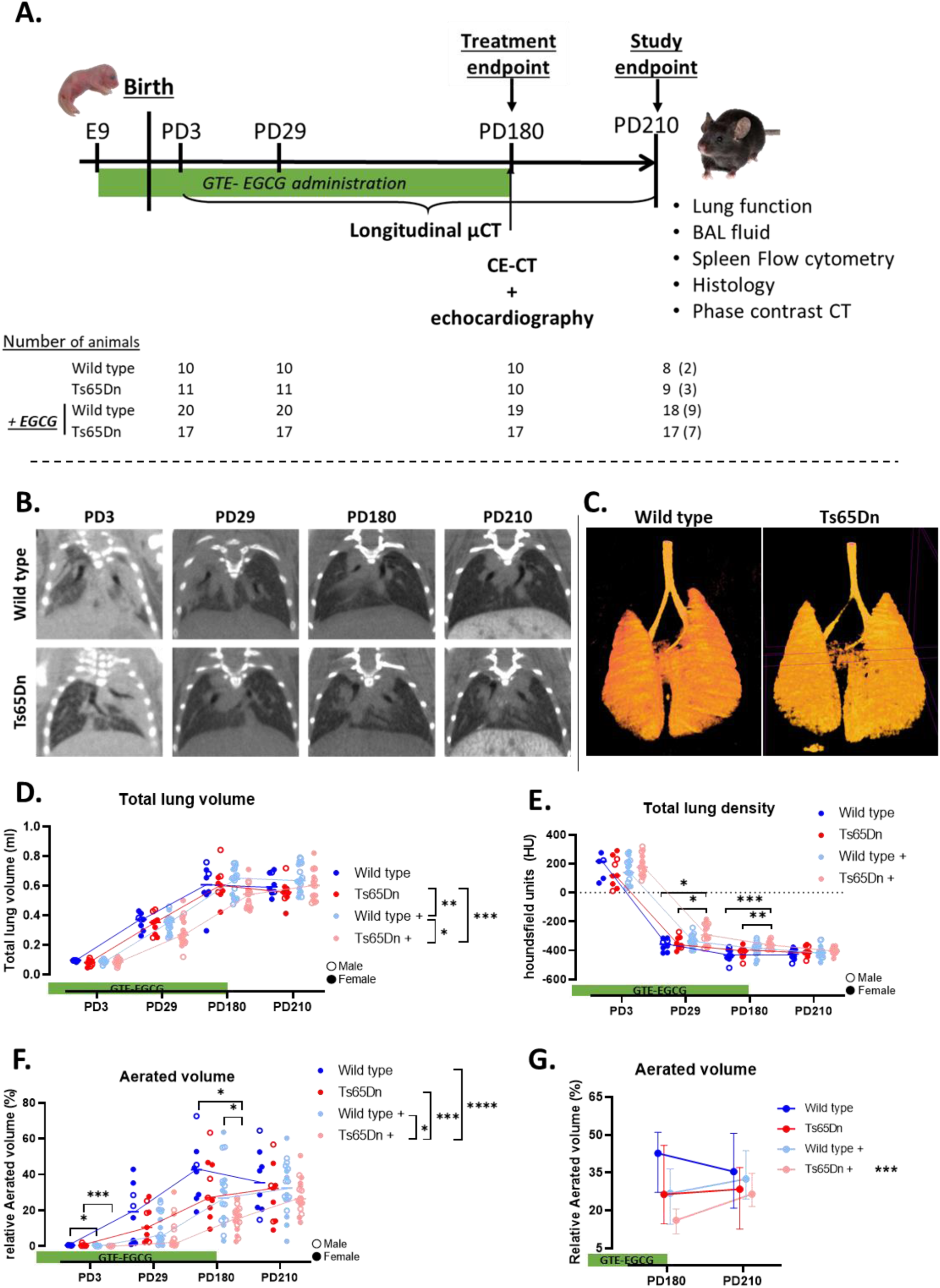
Schematic overview of experimental design and In vivo visualization and longitudinal quantification of lung development in WT and Ts65Dn. (A) GTE-EGCG was administered via drinking water starting at embryonic day 9 (E9) and until PD180. After birth, the mice were scanned using micro-computed tomography (µCT) at postnatal day (PD) 3, PD29, PD180 and PD210. At treatment endpoint, PD180, a contrast enhanced µCT (CE-µCT) and echocardiographic analysis were performed after which GTE-EGCG administration was discontinued. Lung function measurements, immune profile, histological samples and phase-contrast CT were performed at study endpoint (PD210). Four groups were considered which resulted in a number of animals, with the number of males in brackets, after birth. GTE-EGCG = green tea extract – epigallocatechin-3-gallate. (B) Representative images of WT and Ts65Dn mice at PD3, PD29, PD180 and PD210. (C) Three-dimensional (3D) visualization of aerated lung volumes acquired with µCT. µCT-derived biomarkers including; (D) total lung volume in µm^3^, (E) total mean lung density in Hounsfield units (HU) and (F) aerated lung volume corrected according to total lung volume and expressed as percentage (%) were plotted for WT (blue line), Ts65Dn (red line) and GTE-EGCG-treated WT (light blue line) and Ts65Dn (light red line) mice. (G) Detailed analysis of the aerated volumes at treatment endpoint (PD180) and study endpoint (PD210). Data are presented as (D-F) individual values along with group median and (G) median with interquartile range. Males (open dot) and females (full dot) were separated accordingly. * p < 0.05, ** p < 0.01, *** p < 0.001. n = 8-18 per group.

### Non-invasive in vivo µCT shows similar structural lung development in wild type and Ts65Dn mice

First, trisomic and euploid mice were scanned longitudinally with high resolution, low-dose µCT starting at PD3 up to PD210 to track lung development in the different genotypes. We first investigated lung growth by means of total lung volume. Qualitative analysis showed no differences in WT and TS (Figure 1B). Quantitative analysis of lung growth macrostructural lung biomarkers showed a normal lung growth and no significant differences between total lung volume of WT and Ts65Dn mice throughout development (Figure 1D). Then, we evaluated lung maturation by means of total lung densities. The qualitative analysis showed blackening of the lungs throughout lung development (Figure 1B), similar to the quantitative analysis which showed a decrease in total lung density (Figure 1E). Next, we evaluated airway maturation and alveolar growth represented by aerated lung volume. The increase in aerated volume over time indicated lung maturation and is comparable to the increase in total lung volume (Figure 1F). A 3D reconstruction of the aerated lung volume (Figure 1C) confirmed a similar lung structure between WT and Ts65Dn mice at experimental endpoint. Comparison of lung development in both genotypes by µCT analysis showed a similar lung growth and lung maturation between WT and Ts65Dn mice, indicated by equal total lung volume, aerated lung volume and total lung densities (Figure 2 D-F).

**Figure 2:**
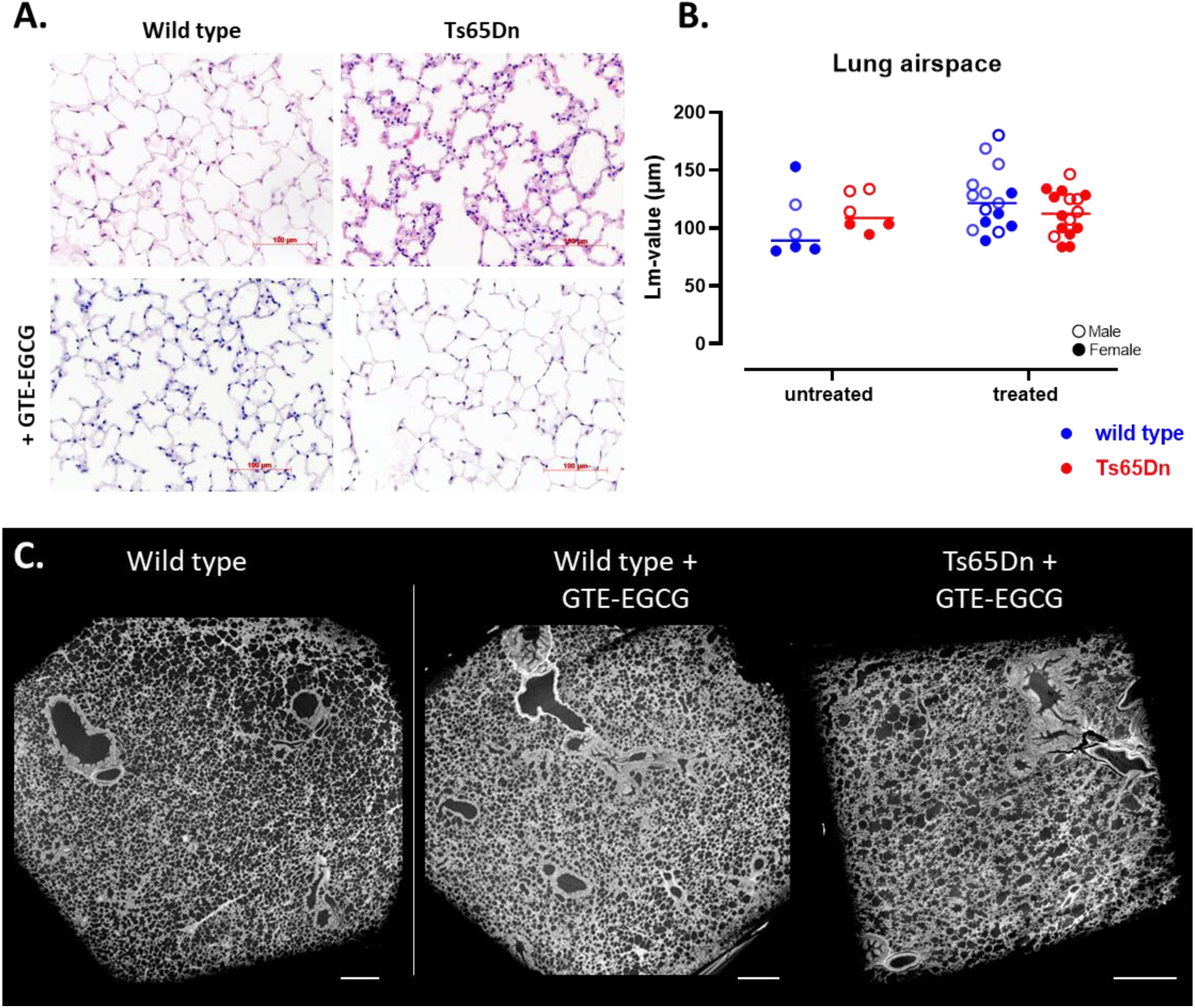
Visualization and microstructural analysis of lung parenchyma. (A). Histological haematoxylin and eosin (H&E) staining (magnification ×20) at study endpoint of WT and Ts65Dn mice with and without GTE-EGCG administration, scale bar 100µm. (B). Lung alveolar airspace (Lm-value) was quantified on histological section of the lung parenchyma represented in (A) and shown for each individual mouse, along with group means. WT (blue) and Ts65Dn (red) with and without treatment of GTE-EGCG, males (open dot) and females (full dot). (C). Phase contrast µCT images showing 3D representation of lung parenchyma (scale bar 250µm with reconstructed voxel size of 2.5 µm).

To get a better insight in the alveolar complexity and alveolar growth, microstructural measurements of airspace enlargement on histological sections of lung parenchyma were performed (Figure 2A). The quantification of airspace enlargement using the mean linear intercepts (Lm) indicated an identical alveolar complexity and suggested a similar lung maturation, thereby confirming similar lung development in WT and Ts65Dn mice (Figure 2B).

Our longitudinal data demonstrated a similar lung development at macrostructural level between WT and Ts65Dn mice. Microstructural analysis at study endpoint using histopathology showed similar lung structural parameters. Hence, both lung macro- and microstructures are alike when comparing WT and Ts65Dn mice.

### Longitudinal µCT biomarkers suggest lung immaturation upon GTE-EGCG administration

After characterizing normal development, we investigated how GTE-EGCG administration modulated the development of the lungs. we assessed the modulatory effect of prenatal GTE-EGCG administration, followed up to six months of age, on macrostructural lung development in euploid and trisomic mice with longitudinal µCT. In both treated groups, GTE-EGCG resulted in decreased lung growth as indicated by a significant decrease in total lung volume throughout development when compared to untreated Ts65Dn mice (Figure 1D). As the effect of GTE-EGCG on total lung volume was significantly more pronounced in Ts65Dn mice than in WT mice, this suggests that Ts65Dn mice presented a larger response to GTE-EGCG (Figure 1D).

We showed a significant increase in total lung density, thereby indicating lower lung capacities in the Ts65Dn treated mice compared to both untreated groups at PD29 (Figure 1E). This difference in total lung density further increased throughout treatment up to treatment endpoint (PD180) confirming a larger response of trisomic mice on GTE-EGCG administration. After discontinuation of GTE-EGCG administration, no variation in mean lung densities were observed at study endpoint thereby suggesting that the effect of GTE-EGCG on lung maturation is transient (Figure 1E).

Next, we analysed aerated lung volume throughout development to investigate lung maturation throughout GTE-EGCG administration. GTE-EGCG administration decreased the aerated lung volume through time, indicating decreased lung capacity in Ts65Dn mice compared to WT treated (p=0.0271), Ts65Dn untreated (p=0.0009) and WT untreated (p<0.0001) groups (Figure 1F). Analysis of each time point separately revealed that GTE-EGCG administration reduced the aerated lung volumes of both WT and Ts65Dn groups compared to the untreated groups right after birth, showing that prenatal administration of GTE-EGCG resulted in early lung immaturation (Figure 1F). The lower aerated lung volume observed at end of treatment between Ts65Dn mice and both WT groups indicated that the effect of GTE-EGCG administration was most pronounced in the Ts65Dn treated mice. However, when GTE-EGCG was discontinued over a washout period of 30 days until study endpoint (PD210), no differences in aerated lung volume were observed on this day (Figure 1F). Similarly, the differences in mean lung densities observed at time of treatment endpoint were no longer present at study endpoint. Additional statistical analysis, comparing aerated lung volume from time of treatment endpoint and study endpoint (Figure 1G), demonstrated a significant difference in the Ts65Dn mice (p=0,0009; 0,1559±0,07 vs 0,2838±0,097), 30 days after GTE-EGCG discontinuation thereby indicating that the effects of GTE-EGCG on lung development and structure were transient and did not persist when GTE-EGCG administration was discontinued.

Microstructural analysis of lung parenchyma using histological analysis at study endpoint, showed a similar alveolar development and lung airspace between WT and Ts65Dn mice (Figure 2A-B). In accordance with and complementary to the µCT-extracted biomarkers at study endpoint, this data confirmed that GTE-EGCG administration had no persistent microstructural effects on alveolar complexity and lung parenchyma. Further 3D microstructural analysis using high-resolution phase contrast CT images of lung parenchyma indicated similar lung maturation (Figure 2C, movies 1-3 in supplementals). Altogether, these data highlight that although the trajectories of pulmonary maturation throughout lung development are different in the context of trisomy and particularly under GTE-EGCG treatment, by adulthood and treatment discontinuation, Ts65Dn and WT mice have overall similar lung airway and parenchymal maturation at study endpoint.

### WT and Ts65Dn littermates present with equal pulmonary lung function and airway reactivity

We performed endpoint lung function measurements to investigate the effect of genotype on pulmonary function. Trisomic and WT mice showed a similar inspiratory capacity (Figure 3A), in line with the aerated lung volumes derived from the µCT data and representing structural biomarkers. Comparison of functional measurements between untreated WT and Ts65Dn mice showed an equally large airway resistance (Figure 3B), tissue elasticity (Figure 3C) and forced vital capacity (FVC) (Figure 3D) indicating that the trisomic mice do not show any restrictive lung disease. Similar to FVC, the Ts65Dn mice demonstrate an equal forced expiratory volume in the first 0,1 second of expiration (FEV_0.1_) as compared to the WT mice (Figure 3E) resulting in equal FEV_0.1_/FVC ratio or “Tiffeneau” index (Figure 3F) and thereby demonstrating the absence of any obstructive phenotype.

**Figure 3:**
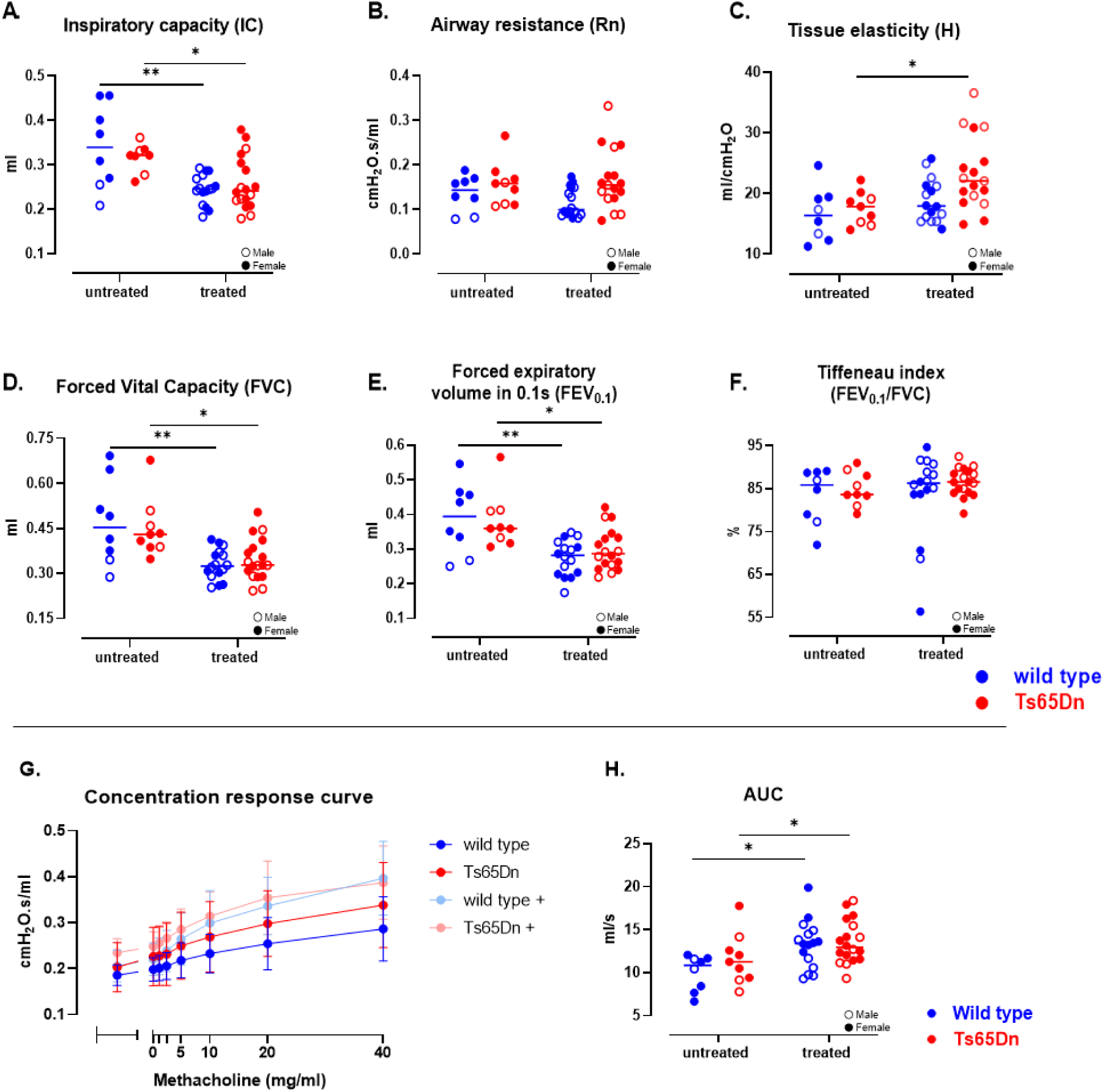
Baseline lung function analysis and airway responsiveness at endpoint in WT and Ts65Dn mice with and without GTE-EGCG modulation. (A) IC was measured using a deep inflation maneuver. FOT respiratory mechanics at baseline showing (B) central airway resistance (R) and (C) tissue elasticity (H). NPFE-derived parameters showing (D) FVC, (E) FEV0.1 and (F) Tiffeneau index (FEV0.1/FVC). (G) Airway resistance was measured in response to increasing methacholine aerosol challenges (0 – 40 mg/ml) in all 4 groups and presented as group mean ± SD. (H) Calculations of the AUC of (G) were presented for all groups. Unless otherwise mentioned, all parameters are shown for each individual mouse, along with group means. WT (blue) and Ts65Dn (red) with and without treatment of GTE-EGCG were shown and males (open dot) and females (full dot) were separated accordingly. * p < 0.05, **p< 0.01. n = 7 – 18 per group.

After the baseline measurements, non-specific airway hyperreactivity was assessed using increasing methacholine concentrations. The Ts65Dn mice showed a similar airway reactivity to inhalation of increasing methacholine concentrations as WT mice (Figure 3G-H).

### GTE-EGCG administration results in declined lung function and airway hyperreactivity in both genotypes

Next, we analysed the lung function data at study endpoint to investigate whether GTE-EGCG administration had an impact on lung function. Both WT and Ts65Dn mice receiving GTE-EGCG up to treatment endpoint (PD180), showed a significantly lower inspiratory capacity at endpoint compared to their corresponding untreated group (Figure 3A). GTE-EGCG administration had no effect on total airway resistance (Figure 3B). However, tissue elasticity was significantly higher in the treated Ts65Dn mice, but not in WT mice (Figure 3C). This indicates a more pronounced effect of GTE-EGCG in trisomic mice compared to euploid mice. Forced expiration measurements demonstrated that GTE-EGCG administration significantly lowered FVC and FEV_0.1_ in both WT and Ts65Dn mice, due to the higher tissue elasticity, whereas Tiffeneau index was not altered between treated and untreated groups (Figure 3D - F). This indicated that the proportional loss of FVC in the first 0.1 second during forced expiration is equal between the groups and we found no signs of an obstructive phenotype in the smaller airways.

Furthermore, we investigated the effect of GTE-EGCG on non-specific airway reactivity using increasing methacholine concentrations. GTE-EGCG administration led to a higher airway reactivity to methacholine and a significant hyperreactive response in the WT mice compared to the untreated WT group (Figure 3G). The treated Ts65Dn mice showed a hyperreactive response observed in relative to the WT untreated, indicated a persistent increased hyperreactive response to GTE-EGCG administration (Figure 3H).

Altogether, we did not observe genotype differences in lung function when comparing untreated euploid and trisomic mice. However, GTE-EGCG results in a persistent decreased lung function and increased airway reactivity in both treated groups 30 days after treatment discontinuation.

To summarize, the overall characterization of Ts65Dn mice regarding structural and functional lung development with longitudinal µCT reveals no genotype differences in lung maturation when compared to their WT littermates up to 7 months follow-up. Similar development of the lung parenchyma was confirmed at the microstructural level by 2D lung histology and 3D PC-µCT. This resulted in equal lung function measures at study endpoint for WT and trisomic mice. However, GTE-EGCG administration resulted in a decrease in lung maturation at treatment endpoint in newborn pups indicated by a lower aerated lung volume, in both genotypes but showing a worse outcome in treated trisomic mice. Although lung maturation and aerated lung volume normalized after GTE-EGCG discontinuation, GTE-EGCG treatment results in persistent decrease in lung function indicated by a lower inspiratory capacity, FVC and FEV_0.1_ in both trisomic and euploid mice. GTE-EGCG administration induced a hyperreactive phenotype in the airways, even after 30 days of treatment discontinuation.

### Characterization of the structural and functional cardiovascular phenotype of Ts65Dn mice and impact of GTE-EGCG modulation

Next to the characterization of the lung, we characterized the pulmonary vasculature and cardiac structure and function by means of a contrast enhanced µCT at PD180 and histology of lung parenchyma for analysis of the pulmonary vasculature. In parallel, we characterized the heart by performing echocardiography at PD180 and histology of the heart at study endpoint.

### Contrast enhanced-µCT analysis in the Ts65Dn mice shows a similar macrovascular structure to WT

Next to characterizing airways and lung parenchyma, we evaluated the pulmonary vascular development driven by the increased risk for developing pulmonary hypertension in the DS population (Bush et al., 2018; Bush and Ivy, 2022). To that end, we subjected WT and trisomic mice to a pre- and post-contrast CE-µCT scan at PD180 to assess *in vivo* macrovascular changes. Visual inspection of the CE-µCT data did not reveal any large structural vascular alterations (Figure 4A-B). Analysis of the CE-µCT-derived biomarkers, demonstrated a similar vascular volume and vascular density between WT and Ts65Dn mice (Figure 4C and 4D). indicating no major macrovascular alterations in the context of trisomy.

**Figure 4:**
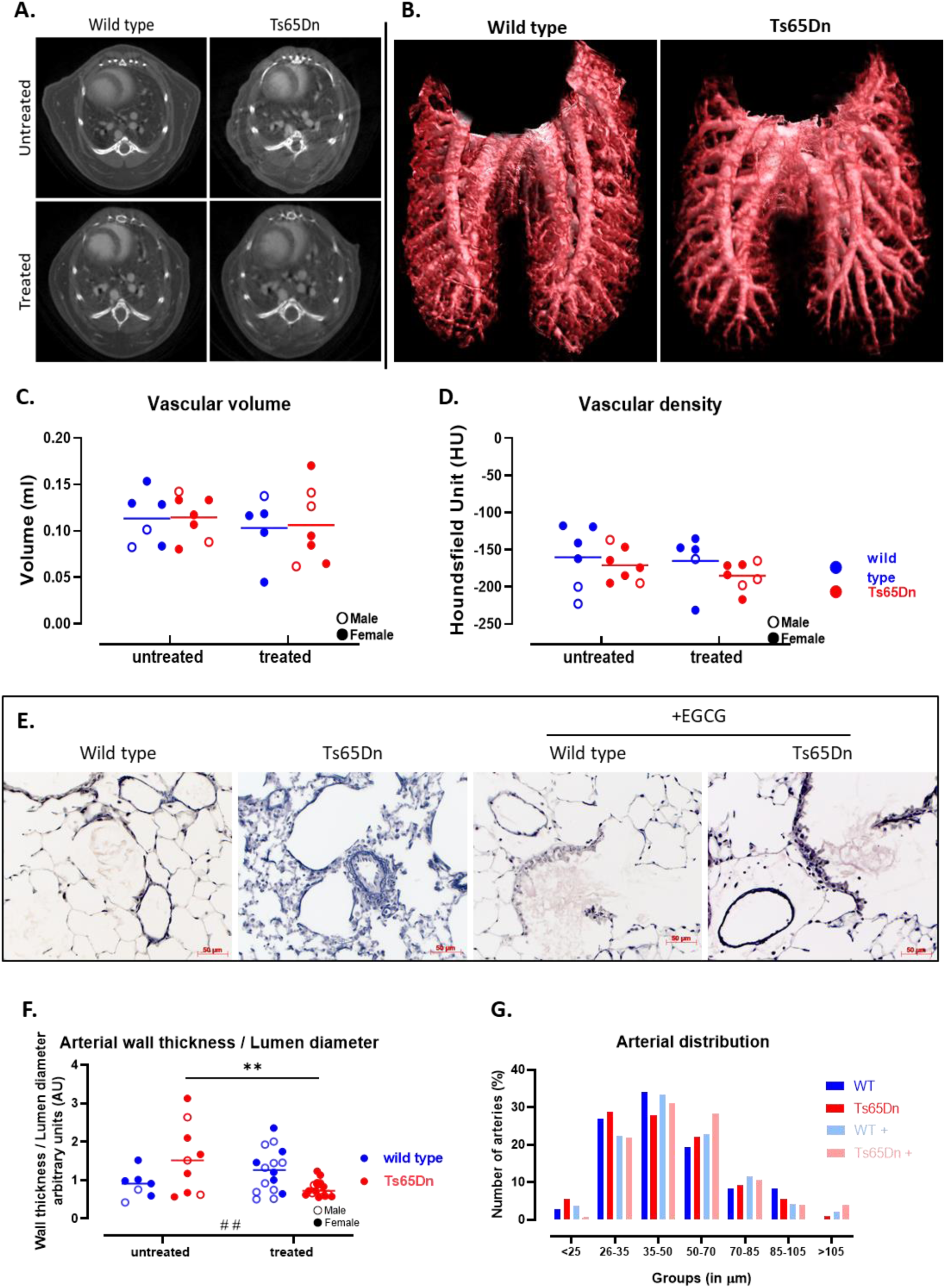
Visualization and quantification of the pulmonary vasculature in WT and Ts65Dn mice using CE-µCT and histology. (A) Representative CE-µCT transversal images of WT and Ts65Dn mice with and without GTE-EGCG administration at treatment endpoint (PD180). (B) 3D reconstructed images of the macrovascular structures in lung ROI of wild type and Ts65Dn mice using the CE-µCT data. (C) Quantification and comparison of vascular volumes and (D) µCT-derived vascular density for wild types and Ts65Dn mice with and without GTE-EGCG treatment (E) Histological representations of small pulmonary arteries (scale bar= 50 µm). (F) Quantification and comparison of arterial wall thickness measured on 12 cross-sectional arteries using histological sections in WT and Ts65Dn mice when administered or not with GTE-EGCG. (G) Graphical representation of the arterial internal diameter grouped according to size and expressed as percentage of total count (n=12). Data are presented as individual values along with group means and males (open dot) and females (full dot) were separated accordingly. * p < 0.05, ** p < 0.01, *** p < 0.001, #, ## indicates a significant interaction between genotype and treatment. n=5-18 per group.

Next, we set out to investigate the vasculature on a microstructural level in order to determine the differences in vascular wall morphology. For this, we performed histological analysis to measure the pulmonary vascular wall thickness (Figure 4E). There was no significant difference in arterial wall thickness or arterial distribution between WT and Ts65Dn mice (Figure 4F and 4G). However, correcting arterial wall thickness to the vascular lumen demonstrated a high variability in the Ts65Dn mice. The Ts65Dn presented a slight increase in wall thickness compared to wild type mice, thereby indicating small signs of vessel hypertrophy in the distal arteries of Ts65Dn mice.

### GTE-EGCG administration decreases arterial wall thickness in Ts65Dn mice

We investigated the effect of GTE-EGCG administration on the vascular component using the CE-µCT reconstructed images. Visual inspection showed a normal macrovascular structure, including size of large pulmonary arteries and similar vascular regions in lung parenchyma, between GTE-EGCG administrated and control groups. The CE-µCT-extracted vascular parameters showed no variation in vascular volume or vascular densities between the 4 groups (Figure 4C-D).

Next, we performed histology of lung parenchyma for detailed analysis of the microvascular level and investigated morphological changes in the microcirculation. GTE-EGCG administration restored the arterial wall thickness and vessel hypertrophy in the Ts65Dn mice to the level of WT mice, thereby indicating that the GTE-EGCG-dependent changes in wall thickness are genotype-specific (Figure 4F).

### RV function is altered in Ts65Dn mice

To evaluate if the slight increase in vessel hypertrophy could lead to cardiac structural and functional changes and to investigate if the Ts65Dn mouse model for DS present any cardiac disorders, we evaluated the cardiac structure and function by performing *in vivo* echocardiographic analysis at treatment endpoint. We first focused on the left ventricle (LV). Calculations of the LV cardiac function showed a similar stroke volume, ejection fraction and cardiac output between WT and Ts65Dn mice, indicating that the trisomic genotype is not associated with an altered LV function (Figure 5A-F). Also, the Ts65Dn mice showed similar left ventricular filling, mitral valve and aortic valve function as the WT, highlighting the absence of any LV alterations. Structurally, estimations of the LV mass were equal in both genotypes. This was confirmed on the CE-µCT images by visual examination at treatment endpoint and by *ex vivo* histological measurements of the heart at study endpoint highlighting similar LV wall thickness (Figure 5K).

**Figure 5:**
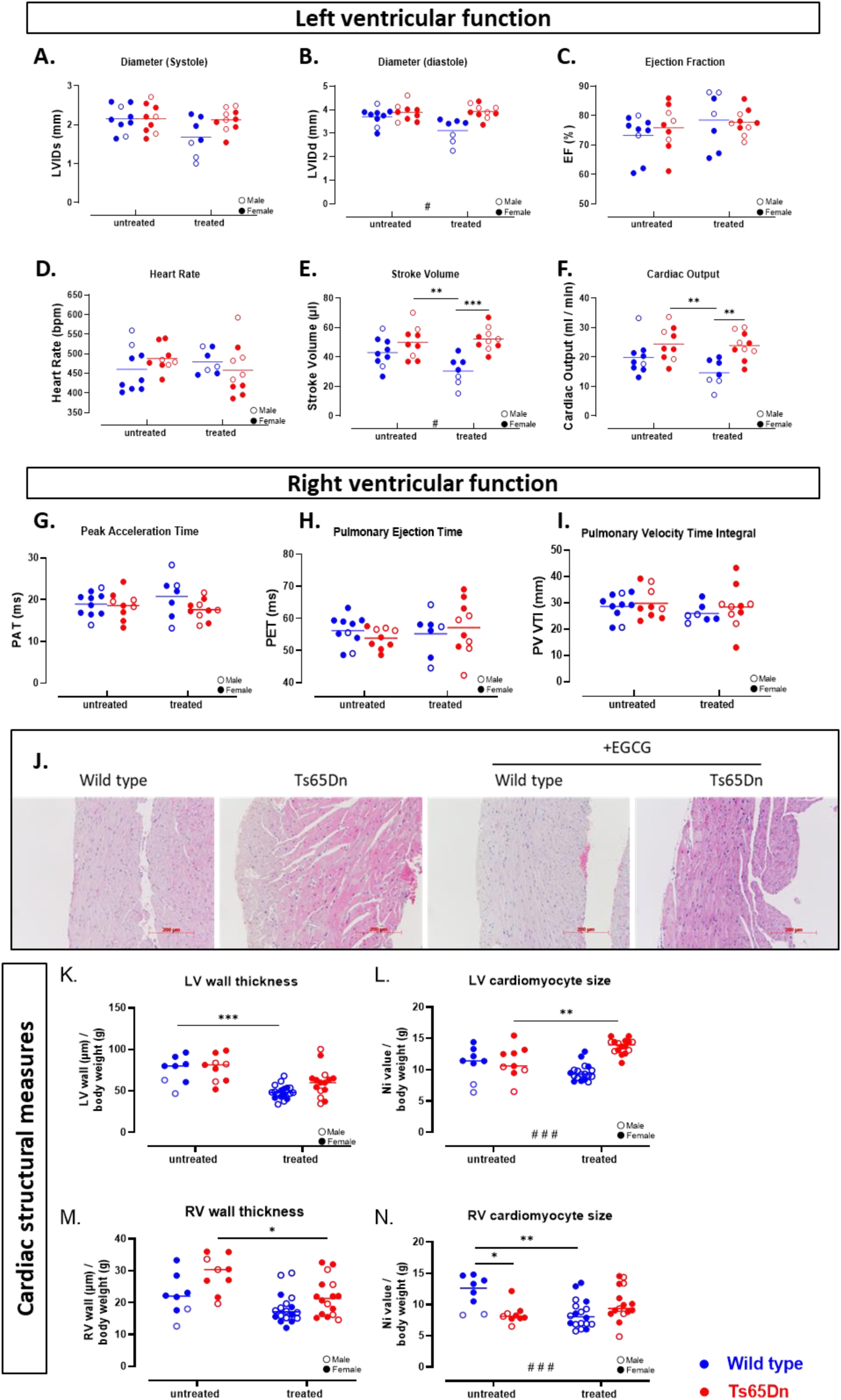
functional and structural analysis of cardiac parameters in in WT and Ts65Dn mice with and without GTE-EGCG modulation. (A-F) Echocardiographic analysis of the LV performed using the PSLAX B-mode resulted in LV inner diameter at end-systole and end-diastole allowing calculated ejection fraction and stroke volume. With corresponding heart rate, cardiac output was measures. (G-I) Using a pulse-wave Doppler Mode from the pulmonary artery using a modified short axis view, pulmonary acceleration time (PAT), pulmonary ejection time (PET) and pulmonary valve velocity time integral for wild types and Ts65Dn mice with and without treatment. J. histological representative figure of the RV wall thickness (scale bar = 200µm). (K-M) *Ex vivo* histological analysis of the heart at study endpoint was performed to measure LV and RV wall thickness, corrected for total body weight. LV and RV cardiomyocyte size were measured counting the number of transections within a reference line. Data are presented as individual values along with group means and males (open dot) and females (full dot) were separated accordingly. * p < 0.05, ** p < 0.01, *** p < 0.001, # indicates an interaction between genotype and treatment. n=6-18 per group.

Next, we investigated the structure of the heart at a microscopic level. Histological quantifications demonstrated similar LV wall thickness in the Ts65Dn mice and the WT mice. In addition, measurement of the individual LV cardiomyocyte size, showed a similar size in the Ts65Dn mice compared to WT (Figure 5L).

Focusing on the right ventricle (RV), echocardiographic analysis of RV function and pulmonary valve function, showed that the Ts65Dn mice have similar right heart functional parameters (including measures as pulmonary acceleration time (PAT), pulmonary ejection time (PET) as indicators of pulmonary hemodynamics and pulmonary vascular resistance and pulmonary valve velocity time integral (PV VTI), an index of stroke volume (Figure 5G-I). At study endpoint, we zoomed in at the microstructural level using *ex vivo* histological analysis, observing that Ts65Dn mice showed a trend towards a thicker RV wall than WT mice (Figure 5M, p=0.0834). Myocardial measures on the number of myocytes, resulted in hypertrophy of the cardiomyocytes in the RV (Figure 5N). These observations are in line with the slight increase in arterial wall thickness and arterial distribution in the Ts65Dn compared to WT mice (Figure 4F-G).

### GTE-EGCG administration affects both LV and RV cardiac development

We investigated the effect of GTE-EGCG administration on the cardiac function using echocardiography at treatment endpoint. First focusing on the LV, our results indicated that GTE-EGCG administration had no effect on LV systolic or diastolic volume, resulting in equal ejection fraction between treated and untreated groups (Figure 5A-C). However, GTE-EGCG administration resulted in a more explicit effect on cardiac functional outcomes in euploid compare to trisomic mice which is seen as a significant genotype difference in stroke volume and cardiac output. Since there is no genotype effect at baseline in the untreated groups, we demonstrate an interaction effect between genotype and GTE-EGCG treatment (Figure 5E and 5F), thereby indicating that the effect of GTE-EGCG administration on cardiac function is genotype dependent.

To look for structural changes, macrostructural analysis of the CE-µCT images was not able to underlie the differences in stroke volume and cardiac output. At microstructural level, endpoint histological measures of the heart showed that GTE-EGCG administration leads to a small dilation of the LV lumen which explains the thinner LV wall in the WT mice (Figure 5K). When calculating the total mass of the LV wall, this appeared to be constant over all the 4 groups.

Next, we focussed on the RV function at treatment endpoint by performing echocardiographic analysis of RV. These functional measures demonstrated that PAT, PET, PAT/PET ratio and PV VTI, indicating pulmonary vascular resistance and an index of stroke volume, showed similar results of RV function between the 4 groups (Figure 5G-I). Furthermore, we demonstrated, using histological analysis of the RV, that GTE-EGCG administration restored the increased RV wall thickness in the Ts65Dn mice up to similar levels as the RV wall thickness of WT mice (Figure 5M). The maximal diameter of the RV (Dmax), increased upon GTE-EGCG administration in both genotypes. However, we did not observe any effect on RV wall thickness or significant effect on number of cardiomyocytes which resulted in a similar size of cardiomyocytes (Figure 5N).

In summary, we characterized the structure and function of the heart cardiopulmonary phenotype of the Ts65Dn mice and indicated no macrostructural changes in pulmonary vasculature with *in vivo* CE-µCT. However, at microvascular level, Ts65Dn mice showed arterial hypertrophy, which is linked with small signs of RV hypertrophy in the Ts65Dn mice which is absent in WT mice. GTE-EGCG administration normalised the arterial thickening observed in Ts65Dn mice and reduced the initiation of RV hypertrophy as indicated by a similar cardiomyocytes size in both treated groups compared to the untreated WT group.

### The systemic and pulmonary immunological status of Ts65Dn mice and effect of GTE-EGCG

#### Ts65Dn mice have less B-cells

To characterize the lung immune status and its potential impact on the lung and vascular phenotype, we assessed the immunological status of Ts65Dn compared to WT mice to reveal potential immunological alterations. Our results regarding pulmonary immunity demonstrated similar numbers of monocytes, lymphocytes, neutrophils and eosinophils in the BAL fluid of WT and Ts65Dn mice (Figure 6A). We investigated the systemic immunological status using flow cytometric analysis of the spleen and showed that Ts65Dn mice have a lower percentage of B-lymphocytes compared to WT mice (Figure 6B). The number of T-lymphocyte and the differentiation into regulatory, cytotoxic and helper T lymphocytes showed similar values in Ts65Dn and WT mice (Figure 6C-F).

**Figure 6:**
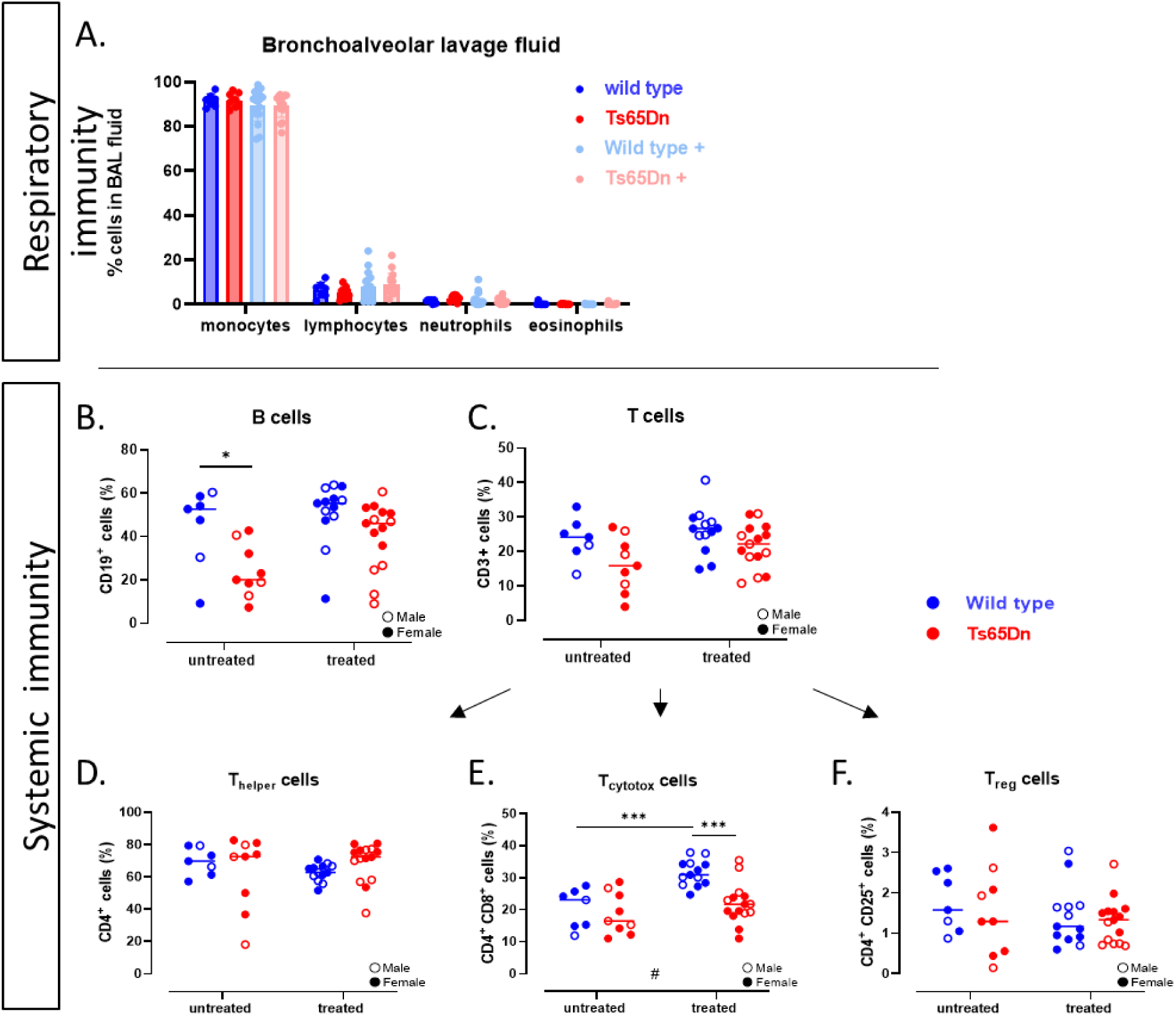
Systemic and pulmonary immunological analysis. (A) Quantification of mono- and polymorphonuclear leukocyte populations in bronchoalveolar lavage fluid showing the relative occurrence of neutrophils (Neutro), lymphocytes (Lympho) and macrophages (Macro). Spleen-digested lymphocytes were stained with a (B) single anti-CD19^+^ (B lymphocytes), (C) anti-CD3^+^ (T lymphocytes), (D) anti-CD3^+^CD4^+^ (Thelper lymphocytes), (E) anti-CD3^+^CD8^+^ (cytotoxic T lymphocytes) and (F) anti-CD3^+^CD4^+^CD25^+^ (regulatory T lymphocytes) expressed as percentage of total cells (B, C) or percentage of total T-lymphocytes (D, E, F). Bars show the mean ± SD. All parameters are shown for each individual value, along with group means. Male (open dot) and female (full dot) mice were separated accordingly. * p < 0.05, ** p < 0.01, *** p < 0.001. n = 7 – 18 per group.

Finally, we investigated the persistent effects of GTE-EGCG administration at PD210 on the immunological phenotype. Pulmonary immunological analysis using BAL fluid cell count showed no persistent effects of GTE-EGCG administration in number of neutrophils, lymphocytes and macrophages between WT and Ts65Dn mice indicating the absence of any infections or potential auto-immune response (Figure 6A). GTE-EGCG administration did not affect total B-lymphocyte number in WT mice, however in the Ts65Dn mice, 30 days after discontinuation of GTE-EGCG the B-lymphocyte population tended to be normalized compared to WT untreated (p=0.0613, Figure 6B).

In summary, we showed a genotype difference with a significantly lower number of B-cells and slightly lower T-cells in the Ts65Dn mice with both being rescued by GTE-EGCG administration. In addition, we showed that GTE-EGCG administration modulates the immune constitution towards a persistent and increased number of cytotoxic T-lymphocytes in WT but not in Ts65Dn mice (Figure 6E).

## Discussion

Motivated by the fact that Individuals with DS present severe clinical cardiovascular and pulmonary abnormalities and the scarcity of preclinical research focussing on the development of these heart and lung alterations in a DS-specific context, we here address the urgent need for a properly characterized animal model representing the clinical respiratory and cardiac deficits. In this study, we here answer two main questions. First, what the cardiopulmonary and immunological phenotype is, throughout development and in the context of trisomy and second, what the effect of GTE-EGCG supplementation is on these systems and in how far effects persisted 1 month after treatment discontinuation.

### What is the Ts65Dn cardiopulmonary and immunological phenotype and to what extent does it replicates the human pathology?

*In vivo* µCT uniquely allowed us to track and quantify lung development immediately after birth until adulthood. We observed that trisomic mouse lungs have a similar lung growth after birth and throughout development as to WT mice indicating that no spontaneous lung pathology was detected. In line with the µCT macroscopic biomarkers, lung histopathology as well as in depth 3D microscopic analysis using PC-µCT showed no signs of lung hypoplasia or airway malacia, indicating a similar airway and alveolar structural development. In parallel, similar lung function measures were found in Ts65Dn and WT mice. In the human situation, whether or not individuals with DS demonstrate increased susceptibility to develop asthma is under debate since the differences observed in lung function in the DS population may be due to the inability to accurately perform lung function testing (Salgueirinho et al., 2016; Weijerman et al., 2011). In the Ts65Dn mice, we observed normal airway reactivity upon increasing methacholine stimulation suggesting that these mice do not present typical characteristics that may be linked to asthma. Also, similar levels of lung eosinophils in both genotypes does not indicate any presence of an asthmatic phenotype in the trisomic mice.

In humans, abnormal pulmonary development is frequently reported in individuals with DS (Bush et al., 2017; Schloo et al., 1991). These anatomical changes include upper airway anomalies and lower airway alterations including decreased alveolar complexity. Most of these observations arise from paediatric studies early in infants and children with DS which demonstrate developmental deficits. Yet, these children with DS have increased frequency of severe lower respiratory tract infections which may impact lung development and induce airway morphological changes (Bloemers et al., 2007; Colvin and Yeager, 2017). We do not observe upper airway alteration or alveolar complexity throughout development, or spontaneous infections. As our mice are housed under semi-sterile conditions in IVC cages, we do not expect exposure to infectious agents. It would be informative whether Ts65Dn mice bear an enhanced risk to infections upon exposure. The increase in respiratory infections observed in clinics is due to the fact that people with DS suffer from an impaired immune response characterized by impaired haematopoiesis (Illouz et al., 2021; Ram and Chinen, 2011; Verstegen et al., 2020). Similar to the clinical situation, the Ts65Dn mice present an immature immune system with decreased B-cells and slightly less T-cells. The effect of recurrent lung infections on lung development of Ts65Dn mice on a DS specific background is yet unknown and may provide more insight in the immunological, functional and structural deficits which are due to the DS genotype.

Besides airway alterations, individuals with DS have an increased risk for significant cardiovascular pathologies including pulmonary hypertension (PH) (Bush and Ivy, 2022). Underlying, this may be due to an increased inflammatory state, lung hypoplasia or the genetic changes in individuals with DS. Infants with DS show increased expression of anti-angiogenic factors resulting in potential abnormal vascular growth in the lung and increasing the risk for developing PH (Galambos et al., 2016).

The slight increase in vascular wall thickness and smaller vascular lumen in the Ts65Dn mice indicates there are small microvascular alterations, potentially making these mice more prone to develop PH in combination with additional stressors. The current hypothesis is that PH initially affects the fragile (pre)capillary bed and small arteries and progressively develops until the large, more proximal vessels show hypertrophy in response to the increase in pulmonary pressures (Humbert et al., 2019). To this regard, our CE-CT data, with extraction of vascular volume and density, showed no changes in the large pulmonary arteries. However, we cannot exclude the presence of vascular pruning and microvascular rarefaction in these Ts65Dn mice. Echocardiographic analysis for structural and functional cardiac changes at an age of 6 months demonstrated equal LV and RV functional parameters. Similar pulmonary pressures on echocardiography suggest that the Ts65Dn mice present a normal vascular development. On the contrary, histopathological examination of the RV implies there is trend towards RV wall thickening in the Ts65Dn mice compared to WT. in addition to the slight increased arterial wall thickness, this may imply that some Ts65Dn pups tend to be more responsive to develop pulmonary vascular complications. We interpret these data as that the cardiac changes are pulmonary-specific and that the RV had to compensate for the hypertrophy observed in the pulmonary arteries to maintain lung homeostasis. Since we do not observe any differences in wall thickness between large and small arteries, we hypothesize that the small changes are rather congenital, indicating small signs of persistent hypertension. Altogether, the increase in RV wall thickness with obvious cardiomyocyte hypertrophy suggests that this effect is acquired in order to restore homeostasis indicating that vascular alterations are present throughout development but do not further progress. These results, showing equal cardiac function but slight alterations in cardiac and vasculature structure, may suggest that these mice might be more prone to develop a PH phenotype.

Other cardiovascular pathologies frequently observed in the DS population is the presence of CHD during birth. The association between DS and CHD is well known, with a reported incidence between 40 and 60% contributing to the morbidity in children with DS (Benhaourech et al., 2016). We do not observe an increased dropout due to severe CHD or another serious congenital dysmorphology in our Ts65Dn mouse population, as we did not observe a clear drop in the number of born pups in trisomic mice.

The immunological deficits we observed are in the same trend compared to the human situation. This is an important finding as it indicates that the Ts65Dn mouse model is suitable to study the effect of lung pathogens from an immunological point of view and thereby investigating the higher incidence of respiratory infections. Furthermore, this model may allow to investigate the impact of additional risk factors of PH, including recurrent infections or toxins on the development of PH in a DS-specific context.

### Is there an effect of prenatal GTE-EGCG administration on heart and lung development?

Although proposed as therapy in individuals with DS to improve cognition, (de la Torre et al., 2016; De la Torre et al., 2014) and ever since heavily studied in the context of its effects on cognition and craniofacial development (Forcano et al., 2021; Starbuck et al., 2021; Wei et al., 2019), this is the first study towards the potential (side) effects of GTE-EGCG on the cardiopulmonary system, nevertheless its known involvement in antiangiogenic and antioxidative pathways (Almatroodi et al., 2020; Chen et al., 2021; Dhatwalia et al., 2018), We here demonstrate that prenatal GTE-EGCG administration has modulatory effects on airway development and cardiac function.

Specifically, we observed that GTE-EGCG administration results in lung immaturation in both WT and Ts65Dn mice evidenced by overall smaller lungs. In addition, µCT biomarkers indicated that airway development is decreased upon GTE-EGCG administration. Discontinuation of GTE-EGCG administration restored lung volume and aerated volumes to the level of the corresponding untreated group, indicating a transient effect of GTE-EGCG on lung immaturation. Although discontinuation of GTE-EGCG administration restored lung structural biomarkers to the level of the corresponding untreated group, the corresponding lung function did remain reduced. We found that GTE-EGCG treatment induced airway hyperreactivity in both WT and trisomic mice.

Interestingly, where trisomic mice had arterial wall thickness with smaller vascular lumen and RV wall hypertrophy, GTE-EGCG administration restored arterial wall thickness and RV wall thickness in the Ts65Dn mice. While no effect on the pulmonary vasculature was observed in the WT group, this highlights a genotype-dependent effect on the pulmonary vascular system. Whether this rescuing effect of GTE-EGCG on the RV is a cause or consequence of GTE-EGCG on the vasculature, remains unclear. Further structural analysis of the microvascular bed and development thereof is required to support our hypothesis that GTE-EGCG administration restores arterial hypertrophy and therefore prevents the development of RV hypertrophy.

Our results suggest that GTE-EGCG administration has a positive effect on restoring the pulmonary vascular system where it seems dysregulated in the context of trisomy. This would be consistent with its angiogenic properties seen in several studies demonstrating the beneficial role of EGCG in cancer prevention by stimulating anti-angiogenic pathways (Dhatwalia et al., 2018; Singh et al., 2011; Zhou et al., 2004). Yet, variations in EGCG concentration or time of administration may also result in differential activation of specific biomolecular pathways. In-dept research towards the pro-or anti-angiogenic signaling pathways during GTE-EGCG administration are necessary to better understand the effect on the pulmonary microvasculature and other organs.

It is well known that recurrent lung infections may affect the pulmonary vascular maturation (Zanoli et al., 2020). Therefore, there effect of GTE-EGCG can impact the vasculature directly or indirectly by rescuing the immune cell maturation. Regarding the effect on immune cells, GTE-EGCG administration rescued the B lymphocyte population in Ts65Dn mice to the level of euploids, suggesting a modulatory effect on immune cells. These results are in line with the modulatory potential of EGCG seen to promote B-cell and T-cell proliferation *in vivo* in a murine leukemia mouse model highlighting the effect of EGCG on the immune response (Huang et al., 2013). In this study, we provide evidence that the Ts65Dn mouse model is suitable to investigate development and effect of therapeutic modulation of respiratory infections to improve the management in a DS-specific population.

Differences in disease models, administration route or concentration and dosage of GTE-EGCG may result in a different outcome and different phenotypes, making it very difficult to compare the effect of EGCG between studies. Administration of EGCG has whole body phenotypic effects which require further research to understand concentration-dependent (side) effects and towards a guideline for safe GTE-EGCG administration in humans (Kim et al., 2014; Younes et al., 2018). This complexity and context-dependent (side) effects of GTE-EGCG administration on different organ systems need further research before its potential treatment potential can be unlocked in a clinical context.

## Conclusion

In the present study, we are the first to fully characterize the cardiopulmonary phenotype of the Ts65Dn mouse model on structural, functional and immunological level. We found a DS-specific phenotype with decreased B-cells and slightly lower number of T-cells, thereby unlocking the relevance and potential of using the Ts65Dn mouse model for respiratory infectious studies in DS. In addition, the Ts65Dn mice present hypertrophy of the RV with increased narrowing of the small blood vessels which suggests these mice are more prone to develop pulmonary vascular diseases suggesting an increased susceptibility to develop PH in combination with additional stressors which is highly relevant to patients with DS that present a high incidence in pulmonary hypertension.

The use of GTE-EGCG administration and the effect on heart and lungs is of important matter since we highlight genotype-independent effects on airway maturation and genotype-specific alterations in the cardiopulmonary system. This study showed that prenatal GTE-EGCG administration results in lung immaturation but shows beneficial effects on the cardiopulmonary system, reducing arterial and RV wall thickness as well as improvements in haematopoiesis with increased number of B-cell lymphocytes. However, we are cautious when formulating potential positive results using GTE-EGCG administration since GTE-EGCG is a freely accessible supplement. The main conclusion of this study is that research regarding the effect of GTE-EGCG supplementation should be considered in a whole-body, holistic approach. we cannot overlook an organ system given the demonstrated pleiotropic response and multi-system nature of DS

## Methods

### Ethics

This study was approved by the institutional animal ethics committee of KU Leuven (P120/2019). All animal experiments were carried out in compliance with national and European regulations. Mice were kept in a conventional animal facility with controlled environmental conditions in individually ventilated cages and free access to food and water.

### Animal model

Euploid and trisomic mice were obtained from our in-house breeding colony at the animal facility of the KU Leuven, established from breeding pairs consisting of wild type (WT) male (003647) and Ts65Dn females (005252, the Jackson Laboratory, Bar Harbor, USA). The date of conception (E0) was determined when a vaginal plug was present. After delivery, the pups were genotyped by PCR from ear snips and allocated in groups according to their genotype, gender and treatment after weaning at postnatal day 21 (PD21).

### Experimental design

Pregnant dams were randomly divided in a non-treatment and treatment-regimen group. After giving birth, the newborn pups were scanned with micro-computed tomography (µCT) at postnatal day (PD) 3, PD29 and at 6 months (PD180), when a contrast-enhanced CT (CE-CT) was performed to study vasculature and heart. After scanning at PD180, treatment endpoint was reached at which treatment was discontinued. At study endpoint (PD210), the mice were scanned using µCT and sacrificed for study endpoint measurements (Figure 1).

### GTE-EGCG treatment

GTE-EGCG was administered to half of the pregnant dams via the drinking water at a concentration of 0.09 mg EGCG/ml. Since EGCG is able to cross the placenta and reach the embryo’s, the treatment was started prenatally at embryonic day 9 (E9) when the lung primordium appears and further develops into trachea and lung buds (Warburton et al., 2010). GTE-EGCG administration continued pre-and postnatally from E9 up to postnatal day 180 (PD180). After weaning at PD21, GTE-EGCG was made available via the drinking water *ad libitum*. The GTE-EGCG solution was prepared daily from a commercially available green tea extract (Mega Green Tea Extract, Life Extension, USA, EGCG content of 45%). Water intake was monitored throughout the experiment.

### Micro-computed tomography (µCT)

At PD3, PD29, PD180 and PD210, mice were scanned using a dedicated *in vivo* micro-CT scanner (Skyscan 1278, Bruker µCT, Kontich, Belgium). Mice were anesthetized by inhalation of 1.5–2% isoflurane in 100% oxygen. The following parameters were used for scanning at PD3 we used 35 kVp X-ray source voltage and 500 μA current combined with a composite X-ray filter of 0,5 mm aluminium and 180 ms exposure time per projection. The scan parameters for mice at PD29, PD180 and PD210 were 55 kVp X-ray source voltage and 700 μA current combined with a composite X-ray filter of 1 mm aluminium, 80 ms exposure time per projection, and overall acquiring projections with 1° increments over a total angle of 360° producing reconstructed 3D data sets with 50 μm isotropic reconstructed voxel size. Software provided by the manufacturer (NRecon, DataViewer, and CTan) was used to reconstruct, visualize, and process μCT data as described previously (Seldeslachts et al., 2022; Vande Velde et al., 2016, 2014). Quantification of the mean lung density, non-aerated lung volume, aerated lung volume, and total lung volume was carried out for a VOI covering the lung, manually delineated on the coronal μCT images, avoiding the heart and main blood vessels. The threshold used to distinguish aerated from non-aerated lung tissue volume was manually set at −493 Hounsfield units (HU) and kept constant over all data sets. Total lung volumes were expressed in mL whereas aerated lung volume was expressed as percentage of total lung volume.

### Contrast enhanced cardiac µCT

At PD180, a pre and post contrast-enhanced µCT **(CE-µCT)** scan was acquired to visualize heart and blood vessels. First, a pre-CE-µCT scan was acquired with similar parameters as the µCT scan except that 20 projections per view were acquired, retrospectively time-based sorted, resulting in a reconstructed 3D data set corresponding to the different phases of the breathing and cardiac cycle (4D) with a scanning time of 12 minutes. Next, a second scan after tail vein injection with a blood pool contrast agent (Exitron nano 12000, Milteny biotec) was acquired which allowed comparison of density changes between pre-and post-contrast scans to extract a biomarker for pulmonary vascular development in the range between fixed thresholds of −345 HU to 385 HU. The data we report here represents the end of the expiratory phase. Reconstruction, analysis and rendering of 3D images was performed using NRecon, CTan, DataViewer and CTvox software (Bruker, Belgium). Adobe Photoshop 2022 has been used to enhance image contrast and overall sharpness.

### Echocardiographic analysis

In parallel with CE-µCT at PD180 (treatment endpoint), echocardiography was performed in mice anesthetized by inhalation of 2% isoflurane. Animals were placed in a supine position on a heating pad to maintain the core body temperature between 37.5–37.7 °C, measured using a rectal probe, to assess the anaesthesia depth and to prevent anaesthesia-induced hypothermia. ECG recordings were performed to monitor the heart and breathing rate. 2-D M-mode echocardiography and tissue and pulsed wave Doppler imaging were performed using a an MS400 (18-38 MHz) transducer connected to a Vevo 2100 echocardiograph (Visualsonics, Canada). LV internal diameters were measured at end-diastole and end-systole (d and s adjuncts, respectively). LV ejection fraction (EF) was calculated as described (Stypmann et al., 2009). In combination with registered heart rate and stroke volume (SV), the cardiac output (CO) was calculated. Left ventricular filling was assessed by pulsed wave Doppler trans-mitral flow tracings (gate size 0.29 mm and Doppler angle − 25°), including E, A, isovolumic contraction time, aortic ejection time, isovolumic relaxation time, and mitral valve deceleration time, just above the tip of the mitral valve leaflets using an apical view. Non-flow time was the sum of isovolumic contraction, aortic ejection, and isovolumic relaxation time. Systolic peak wave, E′, and A′ were measured with tissue Doppler imaging (gate size 0.29 mm and Doppler angle 0°) at the lateral mitral annulus using an apical view. To assess diastolic function, E/A, E/E′, E′/A′, and E/E′/SV ratios were calculated. Using the Pulse Wave Doppler mode, the right ventricle (RV) and pulmonary valve (PV) function were assessed including the pulmonary arterial (PA) acceleration time (PAT), the PA ejection time (PET) and PV velocity-time integral (PV VTI). At least three stable cardiac cycles were averaged for all measurements.

### Pulmonary lung function measurements

Lung function measurements were performed at study endpoint (PD210) using the flexiVent FX system (SCIREQ, EMKA Technologies, Montreal, Canada) equipped with flexiWare 7.6 software and a negative pressure forced expiration (NPFE) FX1 module as previously described (Devos et al., 2017). Briefly, mice were terminally anesthetized with an intraperitoneal injection of pentobarbital (70 mg/kg, Nembutal). Via a tracheotomy, mice were quasi-sinusoidally ventilated with a tidal volume of 10 ml/kg and a frequency of 150 breaths/min to mimic spontaneous breathing. At the start of each measurement, two successive deep inflations were applied to maximally inflate the lungs to a pressure of 30 cm H_2_O in order to open the lungs. Next a deep inflation was performed to determine the inspiratory capacity (IC). Forced oscillation perturbation (FOT) was initiated using the 3 s long, broadband FOT ‘Quick Prime-3’ (QP3) with a frequency between 1 and 20.5 Hz, yielding data on the central airway resistance (Rn), tissue damping (G), tissue elasticity (H) and tissue hysteresivity (eta = G/H). Next, a NPFE (negative pressure-driven forced expiration) perturbation was performed to measure the peak expiratory flow (PEF), forced vital capacity (FVC) and forced expiratory volume in 0.1 s (FEV_0.1_). From these extracted parameters the Tiffeneau index (FEV_0.1_/FVC) was calculated to have an estimate of the obstruction in the smaller airways. Reported values are the average of three accepted measurements for each individual data point. All values were corrected for body weight and presented as such.

Airway hyperreactivity were measured by extracting the airway resistance (Rn)after inhalation of increasing concentrations of methacholine (0, 1.25, 2.5, 5, 10, 20 and 40 mg/mL) After each concentration, the QP3 perturbation was performed 5 times spread over 2 minutes. If the coefficient of determination of a QP3 perturbation was lower than 0.90, the measurement was excluded and not used to calculate the average. Airway resistance (Rn) were plotted against the methacholine concentration and the area under the curve was calculated to perform statistical analysis.

### Immunological analysis

After lung function measures, blood was retro-orbitally collected. 50 µL blood was used for determining whole blood cell counts using the Advia Hematocounter (Siemens Healthineers, Belgium). The rest of the collected blood was centrifuged (14 000 g, 4°C, 10 minutes) and serum was stored at −80°C.

The lungs were lavaged *in situ*, 3 times with 0.7 mL of sterile saline (0.9% NaCl), and the recovered bronchoalveolar lavage (BAL) fluid was pooled. Cells were counted using a Bürker hemocytometer (total cell count) and the BAL fluid was centrifuged (1 000 g, 10 minutes). To obtain the differential cell count, 250 μL of the resuspended cells (100 000 cells/mL) were spun (300 g, 6 minutes) (Cytospin 3; Shandon, TechGen, Zellik, Belgium) onto microscope slides, air-dried and stained (Diff-Quik® method; Medical Diagnostics, Düdingen, Germany). For each sample, 250 cells were counted for the number of macrophages, eosinophils, neutrophils and lymphocytes.

The spleen of each mouse was collected and kept in cold RPMI-1640 medium with Glutamax (Cat: 61870–010, Invitrogen, Belgium). Cell suspensions were obtained by pressing the spleen through a cell strainer (100 mm) (BD Bioscience, Belgium) and rinsing with 10 mL of tissue culture medium. After centrifugation (1 000 g, 10 min), cells were counted on a Bürker hemocytometer and resuspended as 10^7^ cells/mL in complete tissue culture RPMI-1640 medium containing 10% heat-inactivated fetal bovine serum and 10 mg/mL streptomycin/penicillin. If the cell numbers were lower than 10^6^, 100 μL of complete tissue culture medium was added for suspension. After the suspension, 500 000 cells were immunostained with anti-CD3^+^ (APC), anti-CD4^+^ (APC-Cy7), anti-CD8^+^ (PerCP-Cy5.5), and anti-CD25^+^ (PE) or received a single staining with anti-CD19^+^ (PE) labeled antibodies (BD Biosciences, Belgium). Percentages of labeled cells were determined by an LSR Fortessa flow cytometer (BD Biosciences, Belgium). Data was analysed using FlowJo software (TreeStar, Inc, Ashland, OR). Lymphocytes were gated into B cell (CD19^+^), T cell (CD3^+^), T helper cell (CD3^+^, CD4^+^), regulatory T cell (CD3^+^, CD4^+^, CD25^+^), and cytotoxic T cell (CD3^+^, CD8^+^) populations.

### Histopathological and histochemical analysis

During lung harvest, the right lung lobes were clamped, collected and snap frozen in liquid nitrogen and stored at −80° until further analysis. The left lungs were instilled with 4% formaldehyde, inflated at a pressure of 25 cm fluid column and used for histopathological examination. 5 µm sagittal sections were cut and stained with haematoxylin–eosin (H&E) and Sirius red and evaluated by an experienced pathologist in a blinded manner. Airspace enlargement was quantified in 10–15 fields per lung, taken structurally randomized, at a magnification of ×40 by measuring the mean linear intercept (Lm). The pulmonary vascular wall thickness was analysed by measuring the inner and outer diameter of 12 random arteries in each mouse.

Each heart was removed and embedded were immersion-fixed in paraformaldehyde 4%, embedded in paraffin en-block. 5 µm thick sections of transmural tissue were taken from the septal and lateral wall at a mid-ventricular level. Microscopic analysis was performed with a Zeiss Axio Imager ML light microscope (Zeiss, Oberkochen, Germany) measuring septal thickness, LV lumen, maximal diameter of the RV (Dmax) and the lateral wall thickness of the RV and LV at mid-ventricular level. Cardiomyocyte hypertrophy was measured on haematoxylin-eosin (H&E) stained sections, by measuring the Lm value (Dunnill, 1962; Geens et al., 2013). The number of cardiomyocytes transsected by a reference line of 500 µm at x200 magnification was counted on 10 fields per sample and then averaged.

### Phase contrast µCT

Quantitative phase-contrast µCT scans were performed at PETRA III, DESY, Hamburg, Germany at the imaging beamline P05 operated by the Helmholtz-Zentrum Hereon, Geesthacht, Germany. In order to measure differential phase information, a Talbot array illuminator (TAI) was added as wavefront marker to the existing µCT setup. The effective pixel size was at 1.2 µm, using a TAI with a period of 7 µm. Further details about the setup and scan protocol can be found in (Gustschin et al., 2021). The samples were embedded in paraffin wax for scanning, where a beam energy of 20 keV and an exposure time of 65 ms was used. Phase retrieval was computed using the Unified modulated pattern analysis (Zdora et al., 2017). Before tomographic reconstruction using filtered Backprojection, a twofold binning was applied.

### Statistical analyses

The data were analysed using GraphPad Prism (version 9.01. GraphPad Software Inc., San Diego, USA). The data are presented as individual values with mean and standard deviation (SD). For analysis we were interested in the differences between genotype and effect the of treatment. Therefore, we performed a statistical test to answer each of the five research questions. We compared WT and Ts65Dn mice to evaluate if there was a genotype effect; WT and WT treated mice to evaluate if there was a treatment effect in the WT background; Ts65Dn and Ts65Dn treated mice to evaluate if there was a treatment effect in the trisomic background; WT and Ts65Dn treated mice to evaluate if the treatment was rescuing the DS phenotype; WT treated and Ts65Dn treated to evaluate genotype-specific treatment effects. Data of longitudinal µCT-derived biomarkers (mean lung density, total lung volume, non-aerated lung volume and aerated lung volume) was analysed using a mixed-effects model with Geisser–Greenhouse correction. Since time always resulted in a significant difference, as mice grew and their lung volume increased accordingly over time, we focused on differences in the growth patterns as reflected by the interaction effect.

Pairwise comparison between groups at single timepoints were performed as described in (Llambrich et al., 2022). Normality was assessed for each comparison using Shapiro-Wilk and homoscedasticity was assessed using an F-test. Mice identified as outliers by the ROUT test (Motulsky and Brown, 2006) with a Q (maximum desired False Discovery Rate) of 1% were excluded from the analysis. When data were normally distributed, we performed a Welch’s T-test and presented the data as mean and standard deviation. If not, but standard deviations were similar, we performed a Mann-Whitney test and expressed data as median with interquartile range. In case data were not normally distributed and the standard deviations were not similar, we performed a two-sample Kolmogorov Smirnov test. For all tests, statistical significance was assumed when the p value < 0.05. If an interaction between genotype and treatment is present, this is indicated in the graphs with #.

## Acknowledgements

Imaging data was acquired in the Molecular Small Animal Imaging Center (MoSAIC), a core facility of Dept. Imaging and Pathology, Group Biomedical Sciences, KU Leuven. The authors acknowledge the Laboratory Animal Centre core facility of KU Leuven for support with animal care.

## Data availability statement

NA

## Author contributions statement

The study design was developed by BT, SL and GVV. Data collection was performed by BT, SL, LS, JW and EV and data was analysed by BT, JC, HCT, ACJ, FM, MR and BL. Interpretation of the data was done by BT, JH, EV, JV and GVV. Literature search, generation of figures and writing of the manuscript was done by BT and GVV. All authors had final approval of the submitted and published versions.

## Institutional Review Board Statement

The study and animal study protocol were approved by the Institutional Review Board of the Animal Ethics Committee of KU Leuven (ECD number: P120/2019)

## List of Supplementary material online

- Movie1_Lung parenchyma_wild type
- Movie2_Lung parenchyma_wild type treated
- Movie3_lung parenchyma_Ts65Dn treated

